# The evolution of context-specific dominance during selective sweeps

**DOI:** 10.64898/2026.08.12.744435

**Authors:** Carl Mackintosh, Tim Connallon, Filip Ruzicka

## Abstract

Dominance is a widespread feature of genetic variants which affects life-history traits and fitness. Although dominance is generally thought to be an intrinsic property of genetic variants, it can sometimes evolve, as in the classic case of melanism in the peppered moth. The broader question of how likely dominance is to evolve is, however, controversial, because conditions favouring dominance evolution are often restrictive. Here, we revisit Haldane’s classic hypothesis that dominance might evolve during the spread of beneficial mutations to fixation (i.e., during selective sweeps). We first confirm results of earlier models that sweeps of unconditionally beneficial mutations generate little potential for dominance to evolve, even in cases where modifier alleles segregate prior to selective sweeps. However, when sweeping beneficial alleles trade off between different environments — which we explore with the illustrative case of sexually antagonistic selection — the scope for dominance evolution expands. This occurs because modifier alleles can alter dominance separately in each environment, increasing the mean fitness of heterozygotes, prolonging the sojourn time of the sweep, and generating more heterozygosity upon which the modifier can act. In extreme cases, beneficial mutations that were initially destined for fixation can undergo a “dominance reversal” as a result of dominance evolution, converting them to balanced polymorphisms. We quantify how regularly dominance reversals of sweeping sexually antagonistic alleles can be expected to evolve. Overall, our results highlight conditions that allow the dominance of beneficial mutations to evolve, which we discuss in light of data on the frequency of selective sweeps, standing genetic variation for modifiers, and plasticity of modifier effects.

## Introduction

Genetic dominance — the unequal effects of homologous alleles of a gene on traits or fitness —– is a widespread phenomenon with important consequences for population variation (Falconer and Mackay 1996; Charlesworth and Hughes 2000), expression of heritable diseases (Amorim et al. 2017), inbreeding depression (Charlesworth and Willis 2009), and the rate and genetic basis of adaptation (Charlesworth, Coyne, and Barton 1987; Gillespie and Langley 1974; Charlesworth and Charlesworth 2010). Some of the earliest studies in modern genetics (Fisher 1928; Haldane 1939) showed that alleles found at low frequencies are typically recessive relative to common (“wild-type”) alleles (Keightley 1996)—a pattern observed among spontaneous deleterious mutations (Simmons and Crow 1977), genetic variants associated with human diseases (Blekhman et al. 2008), and classical mutant alleles in *Drosophila* and other model organisms (Fisher 1928).

The most widely accepted explanation for the partial recessivity of deleterious mutations is that it is an “intrinsic” property of harmful genetic variants. For example, mutations that decrease the catalytic activity of enzymes can have recessive effects on fitness due to diminishing-returns relationships between enzyme activity and reaction fluxes through metabolic pathways (Wright 1934; Kacser and Burns 1981; Keightley 1996). Intrinsic patterns of dominance can also arise from concave relationships between trait expression and fitness (Manna, Martin, and Lenormand 2011; Sellis et al. 2011; McDonough, Ruzicka, and Connallon 2024), and such models are consistent with data on dominance for deleterious mutations (Charlesworth 1979; Orr 1991; Keightley 1996; Manna, Martin, and Lenormand 2011).

But is this the whole story? While intrinsic mechanisms are likely to explain most observations of dominance, it remains possible that evolution elaborates upon this baseline. Indeed, there is a long-standing alternative tradition of “modifier models” proposing that dominance evolves in response to selection for modifier alleles, situated at other loci, that mediate the fitness of heterozygotes at a focal locus (Fisher 1928; Mayo and Bürger 1997; Billiard, Castric, and Llaurens 2021). Broadly speaking, these models can be put into three classes. The first class of modifier model concerns modifiers that affect alleles at an evolutionarily stable balanced polymorphism (Otto and Bourguet 1999; Spencer and Priest 2016; Llaurens, Billiard, Castric, et al. 2009; Peischl and Bürger 2008; Peischl and Schneider 2010). Here, selection for modifiers can be strong and their evolutionary spread rapid because balancing selection maintains heterozygosity at the target locus, which is the source of selection for modifier alleles. However, balancing selection may only affect a small proportion of functional sites in the genome (reviewed in Ruzicka, Zwoinska, et al. (2025)), restricting the scope for this scenario of dominance evolution.

The second class of modifier model, verbalized by Fisher (1928) and formalised by Wright (1929a), concerns modifiers that reduce the heterozygous expression of rare deleterious alleles maintained at a balance between recurrent mutation and purifying selection (mutation-selection balance). However, the low expected frequency of deleterious alleles limits heterozygosity, such that overall selection for modifiers is weak: on the order of the mutation rate at the locus being modified (Wright 1929a).

The third class of modifier model — and our focus here — proposes that dominance modifiers are favoured during the spread of beneficial alleles in a population (Haldane 1956a) (i.e., during “selective sweeps”; *sensu* Pritchard, Pickrell, and Coop (2010)). Compared to the deleterious mutation scenario, selective sweeps have the advantage of generating extensive heterozygosity at the site of the sweep, with peak heterozygosity occurring when the beneficial allele reaches an intermediate frequency. Furthermore, selective sweeps are common (Messer and Petrov 2013; Corbett-Detig, Hartl, and Sackton 2015; Chen, Glémin, and Lascoux 2020; Castellano, James, and Eyre-Walker 2018; Rousselle et al. 2020; Galtier 2016) and in some cases appear to have favoured the spread of a dominance modifier. For example, the evolution of dominance during a sweep seems to have occurred in the case of industrial melanism, where the dominance of alleles of the ‘melanic’ (dark-pigmented) morph of the peppered moth increased during its spread in industrial regions of England (Haldane 1956a; Kettlewell 1965). Yet so far, theoretical models have not been promising to the idea (Haldane 1956a; Ewens 1966; Wagner and Bürger 1985) because strong selection on the sweeping allele quickly eliminates the heterozygosity that the modifier needs. The reason is nicely summarised by Sved and Mayo (1970): “If high selective values are involved [at the sweep locus], so the intensity of selection for modifiers amongst heterozygotes is high, the gene substitution is completed very quickly. Alternatively, if gene substitution occurs slowly so there are many heterozygotes, the intensity of selection for modifiers must be low”.

One assumption of previous models of dominance evolution during sweeps is that sweeping alleles are unconditionally beneficial. However, it is possible that beneficial mutations undergo trade-offs between contexts of selection during the sweep (e.g., between sexes, different fitness components, or environmental conditions that vary over time and space). Trade-offs can increase the sojourn time and expected heterozygosity of sweeping alleles under net directional selection (Mullon, Pomiankowski, and Reuter 2012; Connallon and Clark 2012). Moreover, it is possible that modifiers might have different effects on dominance in different contexts of selection, as suggested by variation in dominance between environments (e.g., Barson et al. 2015; Karageorgi et al. 2025) and as explored in recent models of sex-specific trait evolution (Spencer and Priest 2016; Siljestam, Rueffler, and Arnqvist 2024; Flintham 2025; Flintham et al. 2026). The theoretical conditions for genetic trade-offs (i.e., discordant phenotypic selection but positively correlated phenotypic effects of genetic variants between environments; Sellis et al. 2011; Connallon and Clark 2014; Moorad and Promislow 2008; Moorad and Hall 2009; Martin and Lenormand 2015) are permissive, with accompanying genomic evidence (Ruzicka, Hill, et al. 2019; Ruzicka, Holman, and Connallon 2022; Rudman et al. 2022; Kinsler, Geiler-Samerotte, and Petrov 2020; Bergland et al. 2014; Fan et al. 2016; Bergland et al. 2014; Johnston et al. 2013; Cole et al. 2024). In particular, in light of growing evidence for dominance reversals (in which alleles are dominant within the context they are favoured; reviewed in (Grieshop, Ho, and Kasimatis 2024)), it seems fruitful to revisit Haldane’s hypothesis for dominance evolution while taking the possibility of trade-offs and dominance reversals into account.

Here, we revisit the potential of selective sweeps to favour the spread of dominance modifiers. Building on earlier models (Haldane 1956a; Sved and Mayo 1970; Wagner and Bürger 1985), we develop analytical approximations for the cumulative selection on unlinked (trans-acting) modifier alleles of arbitrary effect size that increase the dominance of a sweeping mutation. We compare cumulative selection on modifier alleles in two cases: first, an unconditionally beneficial allele, and second, an allele subject to a trade-off — specifically, a sexually antagonistic trade-off — in which the sweeping allele is initially subject to positive selection (so that, in the absence of dominance evolution, the allele is favoured to fix). Modifier alleles can alter dominance in a context-specific manner, such that beneficial reversals of dominance (in which each allele is dominant in the context where it is favoured) can evolve. Finally, we apply our calculations to predict the proportion of completed sweeps that lead to the evolution of dominance reversal at a sexually antagonistic locus through fixation of a dominance modifier allele.

### Model outline

We model a large Wright-Fisher population of *N* diploid individuals with two diallelic loci: a directly selected locus with resident allele *A* and derived allele *a* (arising as a single copy), and a dominance modifier locus with ancestral allele *M* and derived allele *m* (initially segregating at a frequency 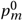). Selection on alleles at the modifier locus can occur due to their effects on dominance at the directly selected locus. We assume selection is weak, random union of gametes, and loose linkage between the two loci such that effects due to linkage disequilibrium are neglected.

We consider two different scenarios where selection favours the derived *a* allele. In the first scenario, *a* is unconditionally beneficial. In the absence of the modifier allele, the genotypes *AA*, *Aa*, and *aa* have relative fitness 1 − *s*, 1 − *hs*, and 1 respectively. The dominance parameter *h* is therefore the dominance coefficient of the *negatively* selected ancestral allele *A*, and modifiers that *decrease h* will be positively selected (this parameterisation is chosen for consistency with the sexual antagonism scenario, where *h*typically represents the dominance of the deleterious effect of a sexually antagonistic allele (Kidwell et al. 1977)). The strength of the modifier effect is given by *ϵ*, which we assume is semi-dominant, such that the dominance coefficient at the selected locus is *h*, *h*(1 *−ϵ*/2) and *h*(1 *−ϵ*) among *MM*, *Mm* and *mm* genotypes, respectively.

In the second scenario, we assume that selection at the *A*locus involves a trade-off. We focus on the illustrative case of sexually antagonistic selection, in which an allele has a fitness advantage in one sex while being detrimental in the other. We suppose that the male-beneficial female-harmful allele is *A*, and the female-beneficial male-harmful allele is *a*. The relative fitness of each genotype in each sex is outlined in Table 1. Without loss of generality, we assume that the fitness cost of expressing the wrong allele is higher in females than in males (*s_f_ > s_m_*), so that the female-beneficial *a* allele is favoured to spread. Further, we assume the dominance modifier allele *m* can have sex-specific effects, altering dominance by a factor 1 − *ϵ_i_* /2 when heterozygous in sex *i*, and by 1 − *ϵ_i_* when homozygous in sex *i* (Table 1). In the absence of the modifier allele, the dominance of each allele is the same in both sexes (*h_f_* = 1 − *h_m_* or “parallel dominance”; Curtsinger, Service, and Prout (1994)), but dominance reversals (in which the antagonistic allele is recessive in the sex in which it is harmful and dominant in the sex in which it is beneficial; or *h_f_ + h_m_ <* 1), can evolve during the sweep.

**Table 1:** Table of relative fitness values when the sweeping allele a is sexually antagonistic.

| **Male** | *AA* | *Aa* | *aa* |
| --- | --- | --- | --- |
| *MM* | 1 | $1 - h_m s_m$ | $1 - s_m$ |
| *Mm* | 1 | $1 - h_m(1 - \epsilon_m/2)s_m$ | $1 - s_m$ |
| *mm* | 1 | $1 - h_m(1 - \epsilon_m)s_m$ | $1 - s_m$ |
| **Female** | *AA* | *Aa* | *aa* |
| *MM* | $1 - s_f$ | $1 - h_f s_f$ | 1 |
| *Mm* | $1 - s_f$ | $1 - h_f(1 - \epsilon_f/2)s_f$ | 1 |
| *mm* | $1 - s_f$ | $1 - h_f(1 - \epsilon_f)s_f$ | 1 |

## Results

### Selection for modifier alleles

#### Unconditionally beneficial sweep

The deterministic trajectories of the sweeping *a* allele and a segregating modifier allele *m* are approximately

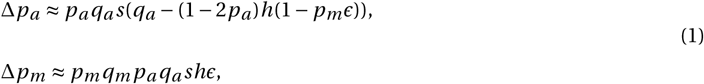

where *q_j_* = 1 − *p_j_* and we neglect terms of order *s*_2_. Selection at the modifier locus is proportional to *p_a_ q_a_*, highlighting the requirement of heterozygosity at the *A* locus. We compare selection for modifier alleles between the two scenarios by calculating “cumulative selection” on *m* across the sweep, given by 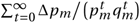. Numerical values for cumulative selection are difficult to interpret, but conclusions can be drawn by comparing values across different regimes. Cumulative selection is proportional to the cumulative heterozygosity of *a* across the sweep, which we approximate by assuming the change in modifier frequency has a negligible effect on the trajectory of the sweeping allele, replacing *p_m_* in Δ*p_a_* with the initial modifier frequency 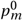. We find cumulative selection on *m* across the sweep of *a* is approximately

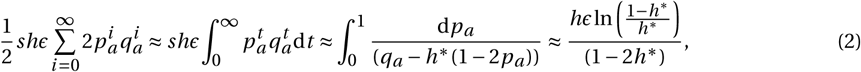

where 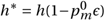. Notably, cumulative selection on the modifier is independent of the strength of selection *s* for the sweeping allele: while higher *s* increases selection for the modifier allele, this effect is cancelled out exactly by the reduction in cumulative heterozygosity during the sweep. As one might expect, cumulative selection on the modifier allele is larger when the sweeping allele is recessive (i.e., the dominance coefficient of the ancestral allele, *h*, is large) and the effect of the modifier (*ϵ*) is large (Figure 1A).

**Figure 1:**
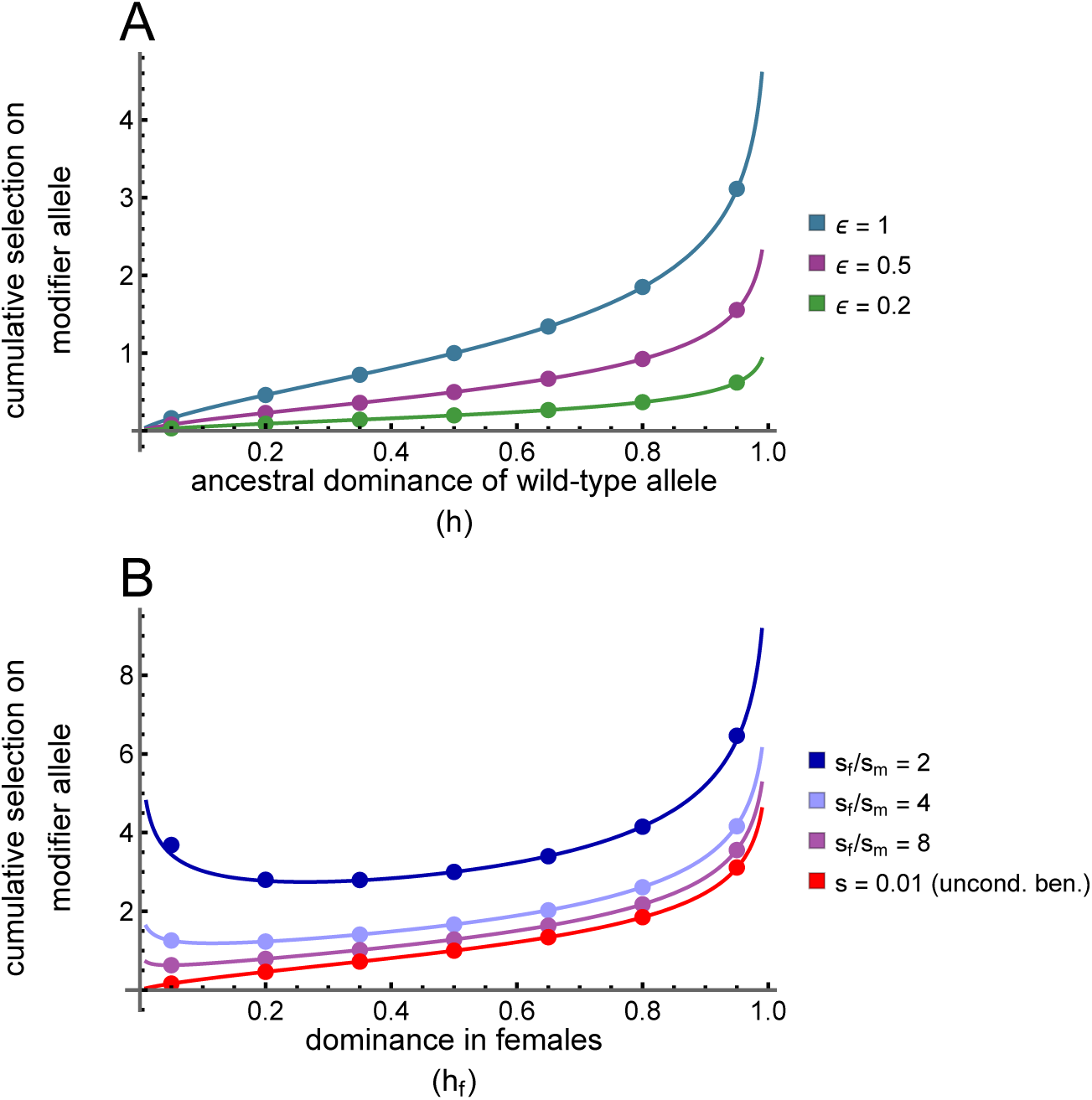
A) Cumulative selection on a dominance modifier allele during a selective sweep of an unconditionally beneficial allele under varying modifier strength; B) cumulative selection in the sexually-antagonistic case under varying degrees of asymmetry in sex-specific selection. Dots show results obtained by iterating exact recursions for 50000 generations (see File S1). For the unconditionally beneficial case, the numerical recursions used *s* = 0.01. In B, *ϵ* = 1, and the mean strength of selection across males and females was fixed at (*s _f_ + s_m_*)/2 *=* 0.01 to match the unconditionally beneficial case. The initial frequency of the modifier allele was 0.0005 (10 copies in a population of size 10000).

#### Sexually antagonistic sweep

When *A* and *a* are sexually antagonistic, the allele frequency change at each locus is

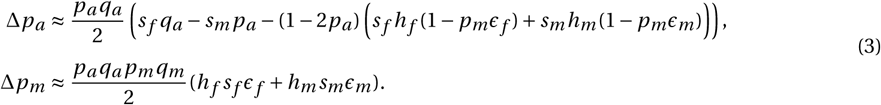

Cumulative selection on *m* across the sweep is now given by

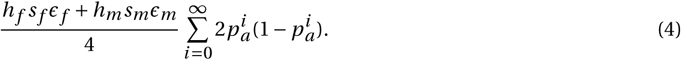

We again approximate a lower bound for cumulative heterozygosity under the assumption that change in frequency of *m* is sufficiently small that it negligibly affects the dynamics of the sweep. The lower bound for cumulative heterozygosity is:

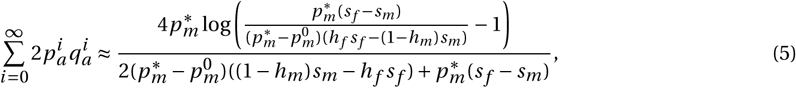

where

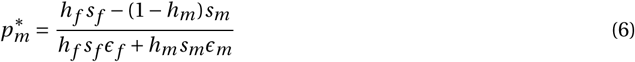

is the critical frequency of *m* above which selection on the sweeping allele transitions from directional to balancing (see Appendix for details). Setting *s_m_* = 0 and replacing *h_f_*, *s*, *ϵ_f_* with *h*, *s*, and *ϵ* recovers the same cumulative selection as for the unconditionally beneficial allele,.

Selection for modifier alleles is particularly high when dominance is initially close to complete in one sex (*h_f_* or *h_m_* close to 0 or 1; Figure 1B). For example, if the female-beneficial *a* allele is initially recessive (i.e., its fitness cost is dominant, so *h_f_* = 1), then directional selection on the sweeping *a* allele will be weak when it is rare, and cumulative heterozygosity will be high. Similarly, if the male-beneficial *A* allele is recessive (i.e., its fitness cost is dominant, so *h_m_* = 1), then directional selection on the sweeping *a* allele will be weak when it nears fixation, and cumulative heterozygosity will also be high (but not as high as when *h_f_* = 1, because we assume *s_f_ > s_m_*). Cumulative selection is also higher with increasing symmetry in sexually antagonistic selection (i.e., the ratio *s_f_* /*s_m_* nears 1). Indeed, when *s_f_ ≈ s_m_*, cumulative selection will be even higher than the approximation suggests, as the modifier significantly increases in frequency during the sweep, thereby altering the trajectory of the sweeping *a* allele (recall that our approximation for the trajectory of *a*assumes no change in the frequency of the *m* allele during the sweep). Even under weak trade-offs between sexes, cumulative selection generated during the sweep is always higher than in the unconditionally beneficial case. Moreover, high symmetry can generate sufficient selection on the modifier allele that its frequency (*p_m_*) often exceeds 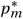 before the sweep is complete. When 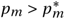, balancing selection at the sweeping locus is generated via a dominance reversal, sustaining heterozygosity at the sweeping locus and leading to fixation of the modifier allele (Figure 2B; C, area left of dashed region but inside solid orange).

**Figure 2:**
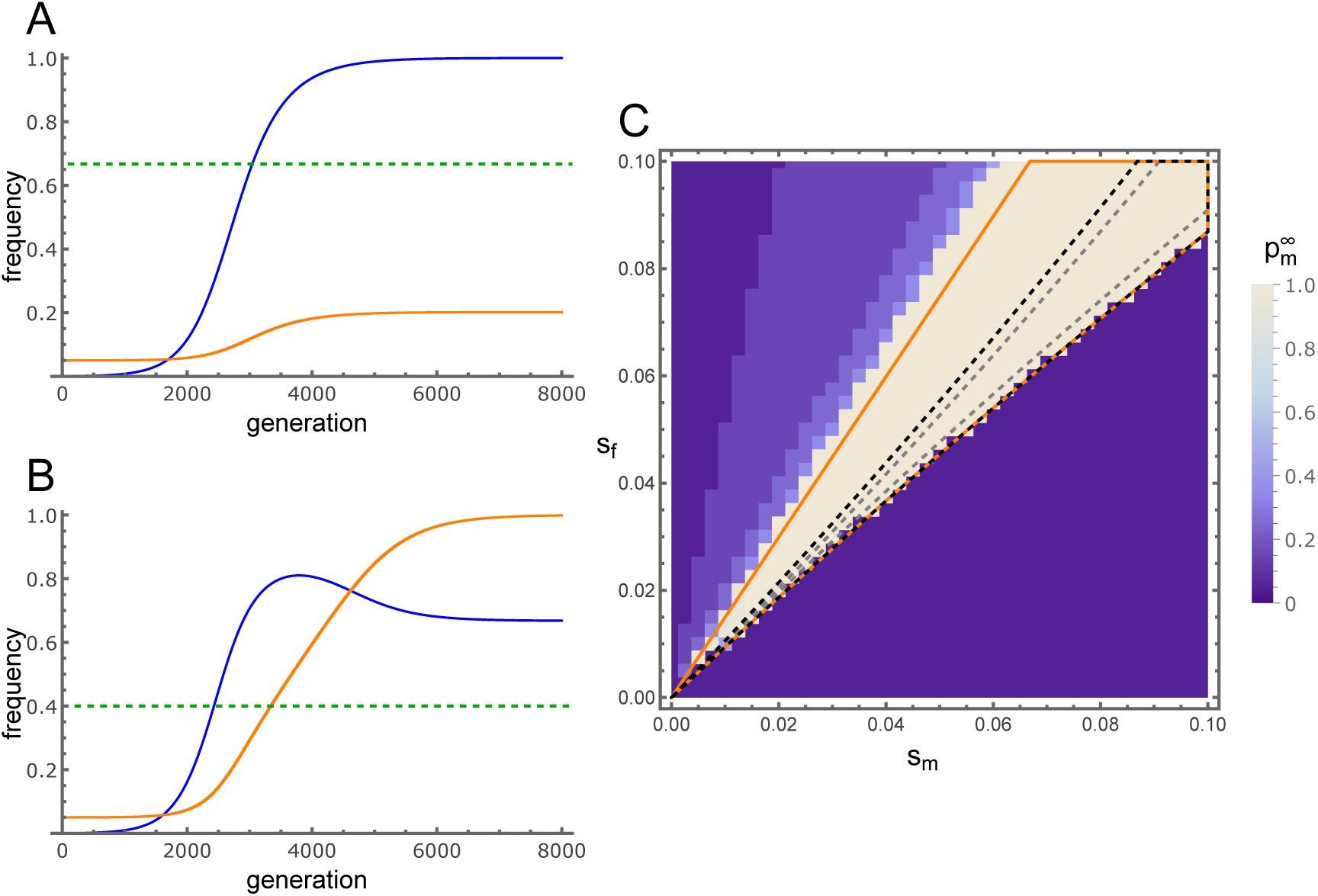
A, B: Deterministic trajectories of the sweeping allele (blue) and the modifier allele (orange) in the sexually antagonistic case. The green dashed line shows the critical modifier allele frequency above which selection on the sweeping allele transitions from directional to balancing. In A, the modifier allele increases in frequency while there is heterozygosity at the sweep locus, whereas in B the modifier allele increase is sufficient to convert the sweep to a balanced polymorphism. C: Final modifier frequency obtain from of forward iteration of the exact recursions that describe the system for 50000 generations. The beige region shows where the modifier allele reached fixation, indicating that the sweep was converted to a balanced polymorphism. The lines show where our analysis predicts balancing selection at the *a* locus. In the absence of the modifier allele, balancing selection occurs within the grey dashed lines. In the presence of the modifier allele (at a constant initial frequency), balancing selection occurs within the black dashed lines. As a result of the increase in frequency of the modifier allele during the sweep (when 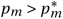), balancing selection occurs within the orange lines. Parameter values used: A) *s_f_* = 0.02, *s_m_* = 0.01, *ϵ_f_* = 0.5, *ϵ_m_* = 0.5, *h_f_* = 0.5, *h_m_* = 0.5, *r* = 0.5; B) as A but with *ϵ_f_ = ϵ_m_* = 1; C) 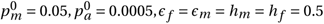.

### Probability a dominance modifier allele converts a sweep to a balanced polymorphism

We have shown that if the *m* allele increases above the critical threshold 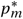 before the *a* allele is fixed, the sweep is converted to a balanced polymorphism through a dominance reversal. What is the probability that this occurs? This depends on two factors: (a) the probability that the modifier allele segregates at the onset of the sweep, and (b) the probability that a segregating modifier allele increases in frequency beyond 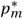 during the sweep.

For a), assuming modifier alleles are derived and selectively neutral in the absence of polymorphism at the target locus, the probability that the modifier locus is *not* polymorphic at the onset of the sweep is

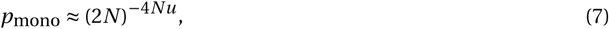

where *u* is the mutation rate to dominance reversal modifier alleles (the approximation is most accurate when 4*Nu ≪* 1; Ewens (2004), Equation 5.68 pp.174).

To find the probability that a modifier allele increases in frequency beyond 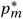 during the sweep (b), we assume that the modifier is segregating as a neutral allele in the absence of polymorphism at the selected locus. Following standard results under neutral theory (Hermisson and Pennings 2005), the distribution of the number of copies of the *m* allele (which we denote as *k*) has probability mass function

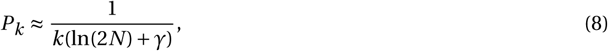

for 1 *≤ k ≤* 2*N −* 1, where *γ ≈* 0.5772 is Euler’s constant. Using Equation 3, the frequency of the modifier allele after the sweep completes, 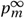, is found by solving the recursive ratio equation

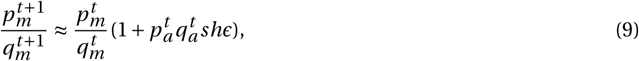

which has the general solution

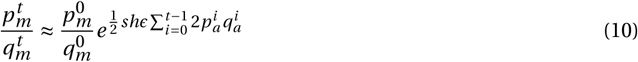

Substituting in Equation 5 shows the modifier frequency at the end of the sweep is

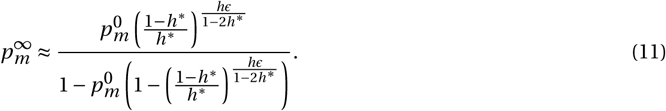

A balanced polymorphism at the directly selected locus occurs if 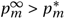. Solving 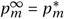 for 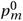 gives the smallest initial frequency of *m* leading to conversion of the sweep, denoted 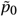. The probability of conversion is therefore the fraction of the initial neutral frequency distribution (Equation 8) above this smallest initial frequency, i.e.: 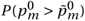. Finally, the overall probability of conversion is 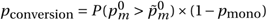

We can then calculate the expected number of completed sweeps it takes for a modifier allele to convert a sweep to a balanced polymorphism. Assuming that combinations of loci experiencing sweeps and their modifiers are independent of other sweep/modifier combinations, this is geometrically distributed with mean 1/*p*_conversion_. If modifier alleles can only evolve and generate dominance reversal at a single locus, many sweeps are predicted to occur before one is converted (Figure 3). If there are *L* loci that could modify dominance in this way, the probability that at least one modifier allele is segregating is 1 *−* (*p*_mono_)*_L_* (which assumes that *Lu* is small enough that the possibility of new modifier alleles arising during the sweep can be neglected). Thus, if 4*NLu* is sufficiently large, modifier alleles appear in the population frequently enough that they segregate with probability approaching 1, in which case the probability that a sweeping allele is converted to a balanced polymorphism becomes 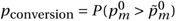.

**Figure 3:**
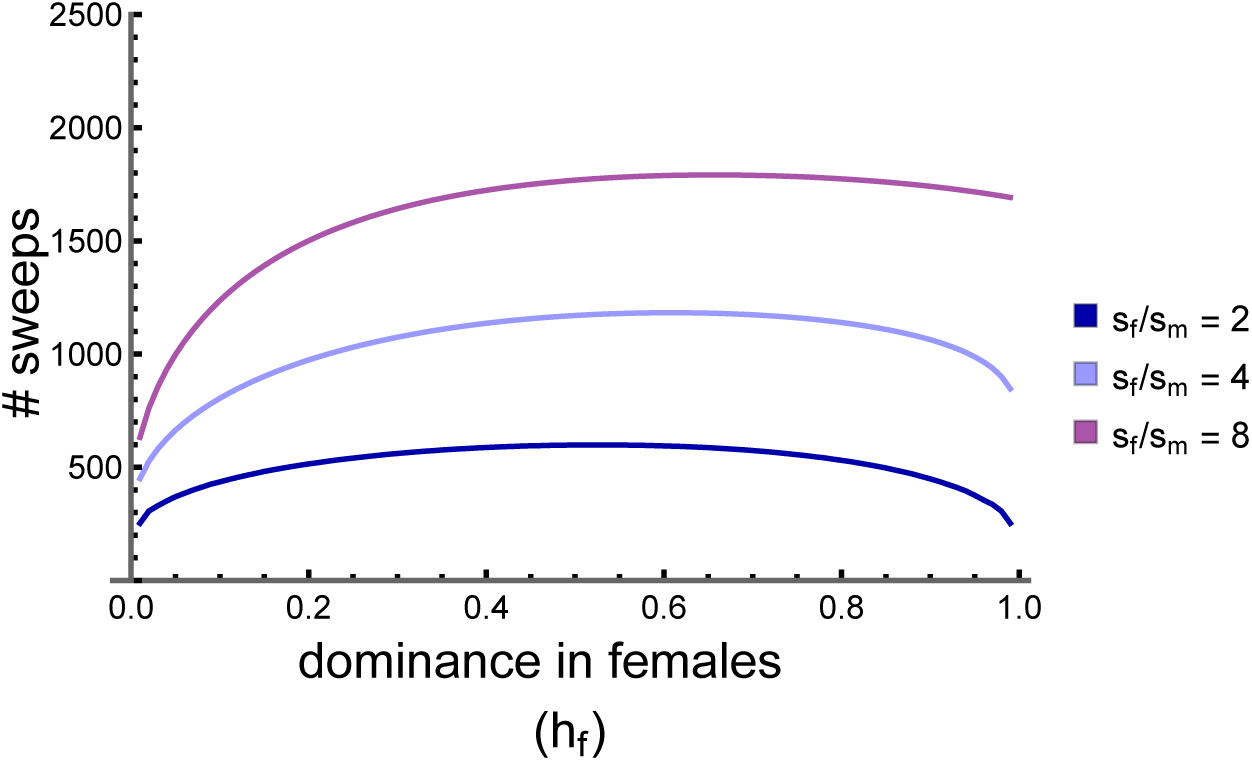
Expected number of completed sweeps of sexually antagonistic alleles required for a dominance reversal modifier allele to convert a sweep to a balanced polymorphism. Modifier alleles generate complete dominance reversals (*ϵ_f_ = ϵ_m_* = 1). Parameters used were *N* = 10^4^, *u* = 10^−8^, (*s_f_ + s_m_*)/2 *=* 0.01.

## Discussion

There are three primary objections to the evolutionary importance of dominance modification, all raised by Sewall Wright nearly a century ago (Wright 1929a; Wright 1929b). The first and most critical objection is that selection for dominance modification requires sufficient heterozygosity at the target locus. In the context of Fisher’s original model where genetic polymorphisms at mutation-selection balance generate the selection for dominance modifiers, selection for dominance modification is extraordinarily weak because alleles at mutation-selection balance generate very little heterozygosity. The second objection is that there may be too few sites in the genome that can mutate to alleles facilitating dominance modification. Taken with the first objection, the evolution of dominance through modifier alleles is hampered if modifier and target alleles do not segregate concurrently. Finally, modifier alleles may have pleiotropic effects on other fitness components (aside from dominance modification), and these direct fitness effects of modifiers might outweigh any indirect effects associated with dominance modification. Wright (1929b) (p. 558) acknowledged a lack of conclusive data regarding the latter two objections, stating that: “Fisher holds on the whole to the affirmative and I am still skeptical, admitting that questions are involved on which we know very little”.

The first objection is mitigated when loci targeted by modifiers segregate for intermediate-frequency alleles (this contrasts with the mutation-selection balance mechanism suggested by Fisher (1928), where heterozygosity is consistently low at each target of dominance modification). Balancing selection, by maintaining intermediate-frequency alleles, can generate sufficiently strong and sustained selection for dominance modification that it can offset direct fitness effects of modifier alleles through pleiotropy (Otto and Bourguet 1999; Peischl and Bürger 2008; Llaurens, Billiard, Leducq, et al. 2008; Peischl and Schneider 2010; Spencer and Priest 2016). Likewise, unconditionally beneficial alleles undergoing selective sweeps each experience a phase where heterozygosity is high and selection for dominance modification can be strong, yet this phase can be fleeting, particularly when selection at the site of the sweep is strong (Haldane 1956b; Sved and Mayo 1970; Wagner and Bürger 1985; this article). A seemingly inescapable conclusion from these models of dominance evolution is that only balanced polymorphisms generate the persistently high heterozygosity required for dominance modifiers to appreciably spread within a population.

Here, we show that cumulative selection for dominance modifier alleles is increased when sweeping alleles exhibit fitness trade-offs. Sweeps of unconditionally beneficial alleles have two opposing effects on the cumulative selection on a modifier allele: strong selection at the site of the sweep increases the maximum strength of selection for the modifier allele, yet it simultaneously decreases cumulative heterozygosity arising during the sweep. These two effects cancel out, leading to weak net selection for the modifier regardless of the strength of selection at the site of the sweep (Sved and Mayo 1970). In contrast, trade-offs between fitness components, sexes, or environments are expected to weaken the net strength of directional selection during a sweep — which increases the cumulative heterozygosity at the site of the sweep — even though selection *within* each fitness context can remain strong (Connallon and Clark 2012; Mullon, Pomiankowski, and Reuter 2012). Selection on a dominance modifier in one or both sexes can thus remain strong, despite effectively weak directional selection at the locus that the modifier targets.

Although cumulative selection for dominance modifier alleles is higher in the presence of trade-offs, the extent to which dominance modification can occur during sweeps depends on several key parameters, including the abundance of selective sweeps, the extent and strength of trade-offs among potentially sweeping alleles, and the extent of genetic variation for modifiers with plastic effects on dominance (i.e., that differ between sexes, fitness components or environments). There is abundant empirical evidence that selective sweeps are common, with high estimated proportions of amino acid substitutions fixed by positive selection (Galtier 2016; Rousselle et al. 2020), and extensive genome-wide signatures of hitchhiking at neutral sites that is at least partially attributable to sweeps (Corbett-Detig, Hartl, and Sackton 2015; Campos, Zhao, and Charlesworth 2017; Castellano, James, and Eyre-Walker 2018; Chen, Glémin, and Lascoux 2020). Trade-offs may also be common, as implied by theory (Martin and Lenormand 2015; Connallon, Czuppon, et al. 2025) and evidence from genome-wide scans for sexually antagonistic genes (Ruzicka, Hill, et al. 2019; Ruzicka and Connallon 2020; Ruzicka, Holman, and Connallon 2022; Cole et al. 2024; Singh, Hasan, and Agrawal 2023), antagonistic pleiotropy between life-history stages (Kinsler, Geiler-Samerotte, and Petrov 2020; Cole et al. 2024), and fitness trade-offs among environments that vary over time or space (Bergland et al. 2014; Rudman et al. 2022; Hereford 2009; Hoban et al. 2016). Although we did not model these other trade-off scenarios, their allele frequency dynamics are similar to the sexually antagonistic case we have modelled (Connallon, Czuppon, et al. 2025), and we therefore expect similar predictions about dominance evolution to apply in these cases.

Far less clear is the extent to which genetic variation for dominance modification segregates in populations, or the potential for different responses to selection for dominance across sexes, environments, or fitness components. This calls back to Wright’s (Wright 1929a; Wright 1929b) second and third objections to Fisher’s original theory of dominance. A century later, we still know almost nothing about the number of sites capable of mutating to alleles that modify dominance at other sites in a genome, nor do we know whether modifier alleles often have pleiotropic effects that limit (and by how much) their responses to selection for dominance modification. What we do know is that populations often harbor genetic variation for dominance, and that effects of dominance can differ between contexts of selection. Evidence of genetic variation for dominance has been reviewed several times and includes examples of evolutionary responses of dominance to artificial selection, and epistatic effects of genetic background on the dominance relations between alleles at individual loci (for examples, see: Sved and Mayo (1970), Mayo and Bürger (1997), Otto and Bourguet (1999), Bagheri (2006), Billiard and Castric (2011), and Li and Bank (2024)). It is unclear whether these cases of genetic variation for dominance might also show plastic effects that vary across environments, sexes, or life stages, and future work is needed to broadly evaluate this possibility (e.g., through sex-specific artificial selection for dominance modification). At present, there is some evidence that dominance can differ across environments and sexes (including dominance reversals, e.g.: Barson et al. (2015) and Karageorgi et al. (2025), reviewed in Grieshop, Ho, and Kasimatis (2024)), but not how common such phenomena are.

Our models are consistent with the notion that dominance modification during a selective sweep is all but impossible unless modifier alleles are already segregating prior to the sweep and such modifiers negligibly affect other fitness components. In the context of dominance reversals, thousands of completed selective sweeps might need to occur in plausibly-sized populations before an appropriate dominance modifier allele fixes (Figure 3). Although our model is couched in a two-locus framework, our “modifier locus” can be more liberally viewed as a set of potentially many sites that can modify dominance of a single beneficial allele. Depending on the number of sites in the set, the chance that at least one segregates for modifier alleles prior to the sweep might not be very small. Indeed, our results suggest this must be the case in order for dominance reversals to evolve with any regularity, implying that dominance evolution is much more likely to occur through *trans-* than *cis*-acting modifiers because the latter have smaller mutational targets (Flintham 2025). Even if modifiers have large enough mutational target sizes that they segregate prior to the onset of the sweep, modifiers may initially be under some form of selective constraint (Wright 1929a; Signor and Nuzhdin 2015) which will tend to reduce probabilities of segregation. Finally, there may be other types of modifier (e.g. context-specific expression modifiers) more capable of resolving the conflict (Flintham et al. 2026). Without data on the number of sites that can modify the dominance of a beneficial allele and the extent to which modifiers also respond to direct selection through pleiotropy, it is difficult to quantitatively evaluate how often dominance is likely to evolve. Our study outlines conditions — notably fitness trade-offs and plastic dominance modification between components of the trade-off — that strengthen selection for dominance modification, yet extensive uncertainty remains about the evolutionary potential for a response to this selection. Resolving this uncertainty is ultimately an empirical task.

## Data availability

File S1 is a Mathematica file containing derivations, results of forward iteration, and code used to make figures; also available as a pdf (File S2). Both files will be uploaded to a public repository.

## Acknowledgements

CM is funded by a grant from the Gordon and Betty Moore Foundation (GBMF11489). FR is funded by a H2020 Marie Skłodowska-Curie COFUND fellowship (No. 101034413). A visit by CM to TC at Monash University was made possible by an Australia Partnering award from the BBSRC (award BB/T019921/1 to Max Reuter and TC). We thank the ABiMS computing cluster at Station Biologique de Roscoff for computing time, and Ewan Flintham for insightful discussions.

## Appendix: conditions for polymorphism under sexual antagonism

One possible outcome is that the increase in the modifier frequency alters the effective dominance of the *a* allele such that the sweep is converted to a balanced polymorphism. In the absence of the modifier allele, polymorphism conditions at the *A* locus are

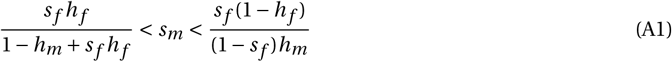

(Kidwell et al. 1977). In the presence of the modifier, conditions for polymorphism are somewhat broader (see Figure 2C, dashed region). Specifically, the conditions for polymorphism are equivalent to (A1), where *h_i_* is replaced with effective dominance coefficients 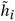:

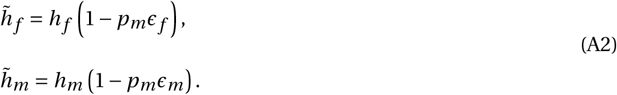

Rearranging for *p_m_* gives the frequency required for the modifier allele to prevent fixation of the sweeping sexually antagonistic allele.

