## Supplementary File 2 for "The evolution of context-specific dominance during selective sweeps"

### Initialisation

```
In[1]:= SetOptions[EvaluationNotebook[], CellContext → Notebook];
        Charting`$InteractiveHighlighting = False;
        SetDirectory[NotebookDirectory[]];
        fixedLabel[text_] := Pane[text, {150, Automatic}]
```

### Unconditionally beneficial allele

#### Analytical results

Mean fitness $\overline{w}$ is given by

```
In[5]:= wbar = (1 - s) (1 - pa)^2 + pa^2 + 2 pa (1 - pa) ((1 - pm)^2 (1 - h s) +
             2 pm (1 - pm) (1 - s h (1 - ϵ / 2)) + pm^2 (1 - h s (1 - ϵ))) // FullSimplify
```

```
Out[5]= 1 - (-1 + pa) s (-1 + pa + 2 h pa (-1 + pm ϵ))
```

The change in the frequency of a is approximately $\frac{p_a(1-p_a)}{2}\frac{\partial \log \overline{w}}{\partial p_a}$:

```
In[6]:= pa (1 - pa) D[Log[wbar], pa] / 2 // FullSimplify;
        % - ((1 - pa) pa s (1 - pa - h (1 - 2 pa) (1 - pm ϵ)))/wbar // FullSimplify
        delpa = ((1 - pa) pa s (1 - pa - h (1 - 2 pa) (1 - pm ϵ)))/wbar;
```

```
Out[7]= 0
```

Assuming weak selection, this is approximately

```
In[9]:= Simplify[Normal[Series[delpa, {s, 0, 1}]]]
```

```
Out[9]= (-1 + pa) pa s (-1 + pa + h (-1 + 2 pa) (-1 + pm ϵ))
```

Similarly, the change in frequency of m is approximately

```
In[10]:= pm (1 - pm) D[Log[wbar], pm] / 2 // FullSimplify;
         % - (h (1 - pa) pa (1 - pm) pm s ϵ)/wbar // Simplify
         delpm = (h (1 - pa) pa (1 - pm) pm s ϵ)/wbar;
```

```
Out[11]=
         0
```

Which again, under weak selection is

```
In[13]:= Simplify[Normal[Series[delpm, {s, 0, 1}]]]
```

```
Out[13]=
         h (-1 + pa) pa (-1 + pm) pm s ϵ
```

Assuming the change in modifier frequency has a negligible effect on the sweep, we can replace $p_m$ in the change in $p_a$ with the initial frequency $p_m^0$.

For a large population, $\Delta p_a \approx \frac{d p_a}{d t}$ and $\sum_{t=0}^{\infty} p_a^t q_a^t \approx \int_0^\infty p_a^t q_a^t \, d t$.

So, cumulative heterozygosity at the sweep locus during the sweep is:

```
In[14]:= Simplify[ Integrate[2 / (s (1 - pa - h (1 - 2 pa) (1 - pm0 ϵ))), {pa, 0, 1},
    Assumptions → Element[h, Reals] && Element[pm0, Reals] && Element[s, Reals] &&
      Element[ϵ, Reals] 0 ≤ pm0 ≤ 1 && 0 < ϵ ≤ 1 && 0 ≤ h ≤ 1] /. h → hstar / (1 - pm0 ϵ)]
```

Out[14]=

$$-\frac{2\,\mathrm{Log}\left[-\frac{\mathrm{hstar}}{-1+\mathrm{hstar}}\right]}{\mathrm{s}-2\,\mathrm{hstar}\,\mathrm{s}}$$

In[15]:= $\mathrm{totHetUB} = \frac{2\,\mathrm{Log}\left[\frac{1-\mathrm{hstar}}{\mathrm{hstar}}\right]}{\mathrm{s}\,(1-2\,\mathrm{hstar})};$

So that cumulative selection on the modifier allele is

```
In[16]:= cumSelUB = s h ϵ totHetUB / 2 /. hstar → h (1 - pm0 ϵ) // Simplify
```

Out[16]=

$$\frac{\mathrm{h}\,\epsilon\,\mathrm{Log}\left[\frac{1-\mathrm{h}+\mathrm{h}\,\mathrm{pm0}\,\epsilon}{\mathrm{h}-\mathrm{h}\,\mathrm{pm0}\,\epsilon}\right]}{1+2\,\mathrm{h}\,(-1+\mathrm{pm0}\,\epsilon)}$$

the recurrence ratio relation $p_m / q_m$ is then approximately

```
In[17]:= (pm + delpm) / (1 - pm - delpm) // FullSimplify;
Series[%, {s, 0, 1}] // Normal;
% // Simplify
```

Out[19]=

$$\frac{\mathrm{pm}\,(-1+\mathrm{h}\,(-1+\mathrm{pa})\,\mathrm{pa}\,\mathrm{s}\,\epsilon)}{-1+\mathrm{pm}}$$

### Numerical recursions

Label the haplotypes as:

MA 1

mA 2

Ma 3

ma 4

The relative fitness of genotype ij is

In[20]:=
```
w11 = 1 - s;
w12 = 1 - s;
w13 = 1 - h s;
w14 = 1 - h (1 - ϵ / 2) s;
w22 = 1 - s;
w23 = 1 - h (1 - ϵ / 2) s;
w24 = 1 - h (1 - ϵ) s;
w33 = 1;
w34 = 1;
w44 = 1;

wbar = w11 x1 ^ 2 + 2 w12 x1 x2 + 2 w13 x1 x3 + 2 w14 x1 x4 +
   w22 x2 x2 + 2 w23 x2 x3 + 2 w24 x2 x4 + w33 x3 x3 + 2 w34 x3 x4 + w44 x4 x4;
```

The following are the genotype frequencies after selection, where xi is the frequency of haplotype i:

In[31]:=
```
p11s = x1 ^ 2 w11 / wbar;
p12s = 2 x1 x2 w12 / wbar ;
p13s = 2 x1 x3 w13 / wbar ;
p14s = 2 x1 x4 w14 / wbar;
p22s = x2 ^ 2 w22 / wbar ;
p23s = 2 x2 x3 w23 / wbar ;
p24s = 2 x2 x4 w24 / wbar ;
p33s = x3 ^ 2 w33 / wbar ;
p34s = 2 x3 x4 w34 / wbar;
p44s = x4 ^ 2 w44 / wbar ;

p11s + p12s + p13s + p14s + p22s + p23s + p24s + p33s + p34s + p44s /. x4 → 1 - x1 - x2 - x3 //
 FullSimplify
```

Out[41]=
```
1
```

Recursions for haplotype frequencies:

In[42]:=
```
x1prime = p11s + (p12s + p13s + p14s (1 - r) + p23s r) / 2;
x2prime = p22s + (p12s + p23s (1 - r) + p24s + p14s r) / 2;
x3prime = p33s + (p13s + p23s (1 - r) + p34s + p14s r) / 2;
x4prime = p44s + (p14s (1 - r) + p24s + p34s + p23s r ) / 2;
```

Check the frequencies sum to 1:

In[46]:=
```
x1prime + x2prime + x3prime + x4prime /. x4 → 1 - x1 - x2 - x3 // FullSimplify
```

Out[46]=
```
1
```

In[47]:=
```
recursions = {x1prime, x2prime, x3prime, x4prime};
```

Convert from haplotype frequencies to allele frequencies and linkage disequilibrium

```
In[48]:= alleleSubEqs = {
           x1 == xA xM + LD,
           x2 == xA (1 - xM) - LD,
           x3 == (1 - xA) xM - LD
          }
         alleleSubs = {x1 → xA xM + LD,
           x2 → xA (1 - xM) - LD,
           x3 → (1 - xA) xM - LD,
           x4 → (1 - xA) (1 - xM) + LD}
Out[48]=
         {x1 == LD + xA xM, x2 == -LD + xA (1 - xM), x3 == -LD + (1 - xA) xM}
Out[49]=
         {x1 → LD + xA xM, x2 → -LD + xA (1 - xM), x3 → -LD + (1 - xA) xM, x4 → LD + (1 - xA) (1 - xM)}

In[50]:= Simplify[Flatten[Solve[alleleSubEqs, {xA, xM, LD}]], x1 + x2 + x3 + x4 == 1]
Out[50]=
         {xA → x1 + x2, xM → x1 + x3, LD → -x1^2 - x2 x3 - x1 (-1 + x2 + x3)}
```

Define functions that iterate the recursions:

```
In[51]:= tSubs = {xA → xA[t], xM → xM[t], LD → LD[t]};

In[52]:= trecs = Simplify[{xA[t + 1] == (x1prime + x2prime /. alleleSubs /. tSubs),
             xM[t + 1] == (x1prime + x3prime /. alleleSubs /. tSubs),
             LD[t + 1] == (-x1prime^2 - x2prime x3prime -
                  x1prime (-1 + x2prime + x3prime) /. alleleSubs /. tSubs)}];

In[53]:= initCons = {xA[0] == 1 - a0,
           xM[0] == 1 - m0,
           LD[0] == 0}
Out[53]=
         {xA[0] == 1 - a0, xM[0] == 1 - m0, LD[0] == 0}
```

Generate a table of allele frequencies for a set of parameters pars, k initial modifier allele copies, and a population of size NN. Iteration for 200000 generations.

```
In[54]:= freqListUB[pars_, k_, NN_] := RecurrenceTable[
           Flatten[{trecs //. pars, initCons /. m0 → k / (2 NN) /. a0 → 1 / (2 NN)}],
           {xA, xM, LD}, {t, 0, 200 000}]
```

Extract $p_m$ and calculate cumulative selection

```
In[55]:= pmListUB[pars_, k_, NN_] := 1 - freqListUB[pars, k, NN][[All, 2]];

In[56]:= cumSelUBNum[pars_, k_, NN_] :=
          With[{list = pmListUB[pars, k, NN]},
           Sum[(list[[i + 1]] - list[[i]]) / (list[[i]] (1 - list[[i]])), {i, 1, Length[list] - 1}]
          ]
```

### Figure 1A

In[57]:=
```
ListPlot[{Table[{H, cumSelUBNum[{s → 0.01, h → H, ϵ → 1, r → 0.5},
       0.0005 * 2 * 10 000, 50 000]}, {H, 0.05, 0.95, 0.15}],
    Table[{H, cumSelUBNum[{s → 0.01, h → H, ϵ → 0.5, r → 0.5},
       0.0005 * 2 * 10 000, 50 000]}, {H, 0.05, 0.95, 0.15}],
    Table[{H, cumSelUBNum[{s → 0.01, h → H, ϵ → 0.2, r → 0.5},
       0.0005 * 2 * 10 000, 50 000]}, {H, 0.05, 0.95, 0.15}]},
   PlotRange → Full, PlotStyle → {{PointSize[0.025], ColorData[1][9]},
     {PointSize[0.025], ColorData[1][6]}, {PointSize[0.025], ColorData[1][8]}}];

Plot[{(1 / 2) s h ϵ (totHetUB) /. hstar → h (1 - pm0 ϵ) /. ϵ → 1 /. pm0 → 0.0005,
    (1 / 2) s h ϵ (totHetUB) /. hstar → h (1 - pm0 ϵ) /. ϵ → 0.5 /. pm0 → 0.0005,
    (1 / 2) s h ϵ (totHetUB) /. hstar → h (1 - pm0 ϵ) /. ϵ → 0.2 /. pm0 → 0.0005},
   {h, 0.01, 0.99}, PlotRange → Full,
   PlotStyle → {ColorData[1][9], ColorData[1][6], ColorData[1][8]}];


plt1A = Legended[
   Labeled[Show[{%, %%}, AxesStyle → Thick, TicksStyle → Directive[Black, 14],
     AxesOrigin → {0, 0}, ImageSize → 350], {Text[Style[
       Column[{"ancestral dominance of wild-type allele", "(h)"}, Center], 16]],
     Rotate[Text[Style[Column[{"cumulative selection on", "modifier allele\n"},
         Center], 16]], π / 2], Text[Style["A", 28]]},
    {Bottom, Left, {Top, Left}}], SwatchLegend[{ColorData[1][9],
     ColorData[1][6], ColorData[1][8]},
    fixedLabel /@ {"ϵ = 1", "ϵ = 0.5", "ϵ = 0.2"}]]
```

Out[59]=

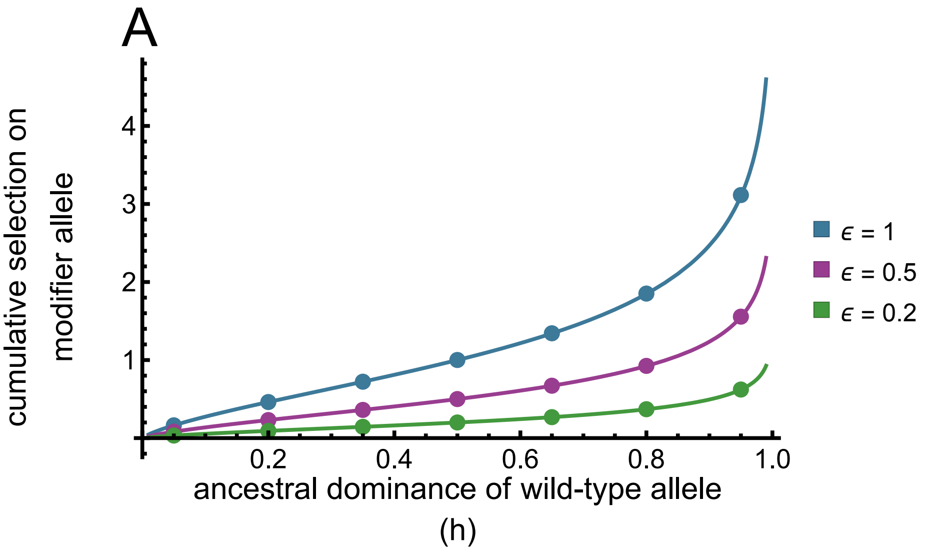

### Sexually antagonistic allele

#### Analytical results

##### Cumulative selection

First, determine the effective dominance coefficient. The relative fitness of each female Aa heterozygote is

In[60]:=
```
wfhetBB = (1 - sf hf);
wfhetBb = 1 - (sf hf) (1 - ϵf / 2);
wfhetbb = 1 - (sf hf) (1 - ϵf);
```

So the mean fitness of female Aa heterozygotes is approximately

In[63]:=
```
meanwfhet = wfhetBB (1 - pm) ^2 + wfhetBb 2 (1 - pm) pm + wfhetbb pm^2 // Simplify
```

Out[63]=
```
1 + hf sf (-1 + pm ϵf)
```

Implying the effective dominance coefficient is

In[64]:=
```
-Coefficient[meanwfhet, sf] // FullSimplify
```

Out[64]=
```
hf - hf pm ϵf
```

Similarly, for males:

In[65]:=
```
wmhetBB = (1 - sm hm);
wmhetBb = 1 - (sm hm) (1 - ϵm / 2);
wmhetbb = 1 - (sm hm) (1 - ϵm);

meanwmhet = wmhetBB (1 - pm) ^2 + wmhetBb 2 (1 - pm) pm + wmhetbb pm^2 // Simplify;
-Coefficient[meanwmhet, sm] // Simplify
```

Out[69]=
```
hm - hm pm ϵm
```

```
effectiveDomSubs = {hf → hf - hf pm ϵf, hm → hm - hm pm ϵm};
```

Mean fitness in each sex and the change in the frequency of a:

In[71]:=
```
wfbar2 = (1 - sf) (1 - pa) ^2 + (1 - hf sf) 2 pa (1 - pa) + pa^2 /.
    effectiveDomSubs // FullSimplify;
D[Normal[Series[Log[%], {sf, 0, 1}]], pa];
wmbar2 = (1 - sm) pa^2 + (1 - hm sm) 2 pa (1 - pa) + (1 - pa) ^2 /.
    effectiveDomSubs // FullSimplify;
D[Normal[Series[Log[%], {sm, 0, 1}]], pa];
% + %%% // Simplify;
delpa2 = (1 / 4) pa (1 - pa) * % // FullSimplify
```

Out[76]=

$$\frac{1}{2}(-1+pa)\,pa\,(sf\,(-1+pa+hf\,(-1+2\,pa)\,(-1+pm\,\epsilon f))+sm\,(pa+hm\,(-1+2\,pa)\,(-1+pm\,\epsilon m)))$$

Show that this can be written in a tidier form:

In[77]:=
```
1/2 (-1 + pa) pa
  (sf (-1 + pa + hf (-1 + 2 pa) (-1 + pm ϵf)) + sm (pa + hm (-1 + 2 pa) (-1 + pm ϵm)));
1/2 (1 - pa) pa
  ( (1 - 2 pa) ( pm (hf sf ϵf + hm sm ϵm) - hf sf - hm sm ) + sf (1 - pa) - sm (pa));
%% - % // Simplify
```

Out[79]=

0

The critical frequency at which selection is no longer directional is given by the smallest value of $p_m$ such that $\Delta\, p_a$ is at some point negative

In[80]:=
```
( (1 - 2 pa) ( pm (hf sf ϵf + hm sm ϵm) - hf sf - hm sm ) + sf (1 - pa) - sm (pa)) /. pa → 1
Solve[% == 0, pm] // FullSimplify
```

Out[80]=

$$hf\,sf-sm+hm\,sm-pm\,(hf\,sf\,\epsilon f+hm\,sm\,\epsilon m)$$

Out[81]=

$$\left\{\left\{pm\to\frac{hf\,sf+(-1+hm)\,sm}{hf\,sf\,\epsilon f+hm\,sm\,\epsilon m}\right\}\right\}$$

Change in the frequency of m

In[82]:=
```
D[Normal[Series[Log[wfbar2], {sf, 0, 1}]], pm];
D[Normal[Series[Log[wmbar2], {sm, 0, 1}]], pm];
delpm = pm (1 - pm) / 4 (% + %%) // Simplify
```

Out[84]=

$$\frac{1}{2}(-1+pa)\,pa\,(-1+pm)\,pm\,(hf\,sf\,\epsilon f+hm\,sm\,\epsilon m)$$

Following a similar argument as before, cumulative heterozygosity is approximately

In[85]:=
```
Integrate[
  1 / ( (1 - 2 pa) ( pm0 (hf sf ϵf + hm sm ϵm) - hf sf - hm sm ) + sf (1 - pa) - sm (pa)),
  {pa, 0, 1}] // Simplify
```

Out[85]=

(Log[sf (-1 + hf - hf pm0 ϵf) + hm sm (1 - pm0 ϵm)] -
Log[sm + hf sf (-1 + pm0 ϵf) + hm sm (-1 + pm0 ϵm)]) /
(sf + sm + 2 hf sf (-1 + pm0 ϵf) + 2 hm sm (-1 + pm0 ϵm)) if condition

Simplifying the numerator, then show it can be written in a tidier form:

In[86]:=
```
Log[(sf (-1 + hf - hf pm ϵf) + hm sm (1 - pm ϵm)) /
    (sm + hf sf (-1 + pm ϵf) + hm sm (-1 + pm ϵm))] /
   (sf + sm + 2 hf sf (-1 + pm ϵf) + 2 hm sm (-1 + pm ϵm)) /. pm → pm0 // FullSimplify;
```

$$\frac{\mathrm{Log}\left[-1+\frac{-sf+sm}{sm+hf\,sf\,(-1+pm0\,\epsilon f)+hm\,sm\,(-1+pm0\,\epsilon m)}\right]}{sf+sm+2\,hf\,sf\,(-1+pm0\,\epsilon f)+2\,hm\,sm\,(-1+pm0\,\epsilon m)};$$

```
% - %% // Simplify
```

Out[88]=

0

Simplify again, where pmcc is the critical modifier frequency:

In[89]:=

$$\frac{\mathrm{Log}\left[-1+\frac{-sf+sm}{sm+hf\,sf\,(-1+pm0\,\epsilon f)+hm\,sm\,(-1+pm0\,\epsilon m)}\right]}{sf+sm+2\,hf\,sf\,(-1+pm0\,\epsilon f)+2\,hm\,sm\,(-1+pm0\,\epsilon m)}$$

$$\frac{pmcc\,\mathrm{Log}\left[\frac{pmcc\,(sf-sm)}{(pmcc-pm0)\,(hf\,sf-(1-hm)\,sm)}-1\right]}{2\,sm\,(1-hm)\,(pmcc-pm0)-2\,hf\,sf\,(pmcc-pm0)+pmcc\,(sf-sm)};$$

$$\%\% - \% \;/.\; pmcc \to \frac{hf\,sf+(-1+hm)\,sm}{hf\,sf\,\epsilon f+hm\,sm\,\epsilon m} \;//\; \mathrm{FullSimplify}$$

Out[89]=

$$\frac{\mathrm{Log}\left[-1+\frac{-sf+sm}{sm+hf\,sf\,(-1+pm0\,\epsilon f)+hm\,sm\,(-1+pm0\,\epsilon m)}\right]}{sf+sm+2\,hf\,sf\,(-1+pm0\,\epsilon f)+2\,hm\,sm\,(-1+pm0\,\epsilon m)}$$

Out[91]=

0

In[92]:=

$$totHetSA = \frac{4\,\mathrm{Log}\left[-1+\frac{-sf+sm}{sm+hf\,sf\,(-1+pm0\,\epsilon f)+hm\,sm\,(-1+pm0\,\epsilon m)}\right]}{sf+sm+2\,hf\,sf\,(-1+pm0\,\epsilon f)+2\,hm\,sm\,(-1+pm0\,\epsilon m)};$$

```
cumSelSA = (sf hf ϵf + sm hm ϵm) totHetSA / 4 // Simplify
```

Out[93]=

$$\frac{(hf\,sf\,\epsilon f+hm\,sm\,\epsilon m)\,\mathrm{Log}\left[-1+\frac{-sf+sm}{sm+hf\,sf\,(-1+pm0\,\epsilon f)+hm\,sm\,(-1+pm0\,\epsilon m)}\right]}{sf+sm+2\,hf\,sf\,(-1+pm0\,\epsilon f)+2\,hm\,sm\,(-1+pm0\,\epsilon m)}$$

#### Determine polymorphism conditions

set up some recursions:

A allele is female beneficial, male deleterious, a allele vice versa.

In[94]:=
```
addSubs = {hf → 1 / 2, hm → 1 / 2};
newFaa = (Maa Faa + Maa FaA / 2 + Maa FAa / 2 + MaA Faa / 2 + MaA FaA / 4 +
     MaA FAa / 4 + MAa Faa / 2 + MAa FaA / 4 + MAa FAa / 4) // Simplify;

newFaA = (Maa FaA / 2 + Maa FAa / 2 + Maa FAA + MaA Faa / 2 + MaA FaA / 2 + MaA FAa / 2 +
       MaA FAA / 2 + MAa FAA / 2 + MAa Faa / 2 + MAa FAa / 2 + MAa FaA / 2 +
       MAA FaA / 2 + MAA FAa / 2 + MAA Faa /. MAa → MaA /. FAa → FaA) // Simplify;

newMAA = (MAA FAA + MAA FAa / 2 + MAA FaA / 2 + MAa FAA / 2 + MAa FAa / 4 +
     MAa FaA / 4 + MaA FAA / 2 + MaA FAa / 4 + MaA FaA / 4) // Simplify;

newMaa = (Maa Faa + Maa FaA / 2 + Maa FAa / 2 + MaA Faa / 2 + MaA FaA / 4 +
     MaA FAa / 4 + MAa Faa / 2 + MAa FaA / 4 + MAa FAa / 4) // Simplify;

newMaA = (Maa FaA / 2 + Maa FAa / 2 + Maa FAA + MaA Faa / 2 + MaA FaA / 2 + MaA FAa / 2 +
       MaA FAA / 2 + MAa FAA / 2 + MAa Faa / 2 + MAa FAa / 2 + MAa FaA / 2 +
       MAA FaA / 2 + MAA FAa / 2 + MAA Faa /. MAa → MaA /. FAa → FaA) // Simplify;

newFAA = ( MAA FAA + MAA FAa / 2 + MAA FaA / 2 + MAa FAA / 2 + MAa FAa / 4 +
     MAa FaA / 4 + MaA FAA / 2 + MaA FAa / 4 + MaA FaA / 4) // Simplify;


Wf = 1 newFaa + (1 - hf sf) newFaA + (1 - sf) newFAA;
Wm = 1 newMAA + (1 - hm sm) newMaA + (1 - sm) newMaa;

Faa′ = newFaa / (2 Wf) /. MAa → MaA /. FAa → FaA;
FaA′ = newFaA (1 - hf sf) / (2 Wf) /. MAa → MaA /. FAa → FaA;
FAA′ = newFAA (1 - sf) / (2 Wf) /. MAa → MaA /. FAa → FaA;
Maa′ = newMaa (1 - sm) / (2 Wm) /. MAa → MaA /. FAa → FaA;
MaA′ = newMaA (1 - hm sm) / (2 Wm) /. MAa → MaA /. FAa → FaA;
MAA′ = newMAA / (2 Wm) /. MAa → MaA /. FAa → FaA;

Faa′ + FaA′ + FAA′ // Simplify
Maa′ + MaA′ + MAA′ // Simplify

recursions = {Faa′, FaA′, FAA′, MAA′, MaA′, Maa′};
```

Out[109]=

$$\frac{1}{2}$$

Out[110]=

$$\frac{1}{2}$$

denote female beneficial allele frequency by paf in females, pm in males. Fx is coefficient of inbreeding in sex x.

In[112]:=

```
subeq = {FAA → (1 - paf) ^2 (1 - Ff) + Ff (1 - paf),
   FaA → 2 paf (1 - paf) (1 - Ff),
   Faa → (paf) ^2 (1 - Ff) + Ff paf,
   MAA → (1 - pam) ^2 (1 - Fm) + Fm (1 - pam),
   MaA → 2 pam (1 - pam) (1 - Fm),
   Maa → (pam) ^2 (1 - Fm) + Fm pam};
{FAA == (1 - paf) ^2 (1 - Ff) + Ff (1 - paf),
  FaA == 2 paf (1 - paf) (1 - Ff),
  Faa + FaA + FAA == 1,
  MAA == (1 - pam) ^2 (1 - Fm) + Fm (1 - pam),
  MaA == 2 pam (1 - pam) (1 - Fm),
  Maa + MaA + MAA == 1};
subs =
 Flatten[Solve[{%[[1]], %[[2]], %[[4]], %[[5]]}, {pam, paf, Fm, Ff}]] // Simplify // Normal
```

Out[114]=

$$\left\{pam \to 1 - \frac{MaA}{2} - MAA,\ paf \to 1 - \frac{FaA}{2} - FAA,\ Fm \to \frac{MaA^2 + 4\,MaA\,MAA + 4\,(-1 + MAA)\,MAA}{MaA^2 + 4\,(-1 + MAA)\,MAA + MaA\,(-2 + 4\,MAA)},\ Ff \to \frac{FaA^2 + 4\,FaA\,FAA + 4\,(-1 + FAA)\,FAA}{FaA^2 + 4\,(-1 + FAA)\,FAA + FaA\,(-2 + 4\,FAA)}\right\}$$

Determine conditions for the 0 and 1 equilibria to be unstable

In[115]:=

```
jacob = Simplify[
   Transpose[{D[recursions, Faa], D[recursions, FaA], D[recursions, FAA],
     D[recursions, MAA], D[recursions, MaA], D[recursions, Maa]}]];
Simplify[jacob /. subeq /. paf → 0 /. pam → 0];
Eigenvalues[%] // FullSimplify
```

Out[117]=

$$\left\{0, 0, 0, 0, 0, \frac{1}{2}\left(\frac{-2 + sf + hf\,sf}{-1 + sf} - hm\,sm\right)\right\}$$

In[118]:=

```
Simplify[jacob /. subeq /. paf → 1 /. pam → 1];
Eigenvalues[%] // FullSimplify
```

Out[119]=

$$\left\{0, 0, 0, 0, 0, \frac{-2 + sm + hm\,sm + hf\,(sf - sf\,sm)}{2\,(-1 + sm)}\right\}$$

In[120]:=

$$eval1 = \frac{-2 + sm + hm\,sm + hf\,(sf - sf\,sm)}{2\,(-1 + sm)};$$

$$eval0 = \frac{1}{2}\left(\frac{-2 + sf + hf\,sf}{-1 + sf} - hm\,sm\right);$$

When both are unstable, the equilibrium allele frequency is intermediate and so we have polymorphism

In[122]:=

```
sm /. Solve[eval1 - 1 == 0, sm][[1]] // FullSimplify
sm /. Solve[eval0 - 1 == 0, sm][[1]] // FullSimplify
polyConditions = (sm > %% && sm < %) // Simplify
```

Out[122]=

$$\frac{hf\ sf}{1 - hm + hf\ sf}$$

Out[123]=

$$\frac{(-1 + hf)\ sf}{hm\ (-1 + sf)}$$

Out[124]=

$$sm > \frac{hf\ sf}{1 - hm + hf\ sf}\ \&\&\ sm < \frac{(-1 + hf)\ sf}{hm\ (-1 + sf)}$$

The modifier allele augments these conditions

In[125]:=

```
polyConditionsMod = polyConditions /. effectiveDomSubs // Simplify
```

Out[125]=

$$sm > \frac{hf\ sf\ (-1 + pm\ \epsilon f)}{-1 + hm + hf\ sf\ (-1 + pm\ \epsilon f) - hm\ pm\ \epsilon m}\ \&\&\ sm < \frac{sf\ (-1 + hf - hf\ pm\ \epsilon f)}{(-1 + sf)\ (hm - hm\ pm\ \epsilon m)}$$

From this, we can again find the critical modifier frequency at which polymorphism is unstable (assuming sf > sm)

In[126]:=

$$\text{Flatten}\left[\text{Solve}\left[\frac{hf\ sf\ (-1 + pm\ \epsilon f)}{-1 + hm + hf\ sf\ (-1 + pm\ \epsilon f) - hm\ pm\ \epsilon m} - sm == 0,\ pm\right],\ 1\right]\ //\ \text{Simplify}$$

```
pmc = pm /. %
```

Out[126]=

$$\left\{pm \to \frac{hf\ sf\ (-1 + sm) + sm - hm\ sm}{hf\ sf\ (-1 + sm)\ \epsilon f - hm\ sm\ \epsilon m}\right\}$$

Out[127]=

$$\frac{hf\ sf\ (-1 + sm) + sm - hm\ sm}{hf\ sf\ (-1 + sm)\ \epsilon f - hm\ sm\ \epsilon m}$$

Which is equivalent to the previous condition under weak selection:

In[128]:=

$$\text{Normal}\left[\text{Series}\left[\frac{hf\ sf\ (-1 + sm) + sm - hm\ sm}{hf\ sf\ (-1 + sm)\ \epsilon f - hm\ sm\ \epsilon m}\ /.\ sf \to sf\ \epsilon\ /.\ sm \to sm\ \epsilon,\ \{\epsilon, 0, 0\}\right]\right]\ //\ \text{FullSimplify}$$

Out[128]=

$$\frac{hf\ sf + (-1 + hm)\ sm}{hf\ sf\ \epsilon f + hm\ sm\ \epsilon m}$$

#### Probability of conversion of sweep to a balanced polymorphism

First, solve the ratio recursion for the final modifier frequency:

In[129]:=

```
finalModFreqSA =
 qt /. Solve[qt / (1 - qt) == pm0 / (1 - pm0) Exp[(sf hf ϵf + sm hm ϵm) / 4 totHetSA] , qt][[
   1]] // FullSimplify
```

Out[129]=

$$1+\frac{-1+\text{pm0}}{1+\text{pm0}\left(-1+\left(-1+\frac{-\text{sf}+\text{sm}}{\text{sm}+\text{hf sf }(-1+\text{pm0 }\epsilon\text{f})+\text{hm sm }(-1+\text{pm0 }\epsilon\text{m})}\right)^{\frac{\text{hf sf }\epsilon\text{f}+\text{hm sm }\epsilon\text{m}}{\text{sf}+\text{sm}+2\text{ hf sf }(-1+\text{pm0 }\epsilon\text{f})+2\text{ hm sm }(-1+\text{pm0 }\epsilon\text{m})}}\right)}$$

Then, find the smallest number of initial copies $k$ such that the modifier allele frequency exceeds the critical frequency, assuming $k$ is initially distributed according to $p_k$. Conditioned on segregation of the modifier allele, the proportion of the distribution above this number is the probability the sweep is converted to a balanced polymorphism.

In[130]:=

$$\text{pk} = \frac{1}{\text{k }(\gamma+\text{Log}[2\text{ NN}])};$$

```
probConversion[params_, pop_] :=
 With[{mink = k /. FindRoot[(finalModFreqSA //. params /. pm0 → k / pop) ==
       (pmc //. params /. pm0 → k / pop), {k, 10}]},

  1 - Sum[pk /. γ → EulerGamma /. NN → pop, {k, 1, Floor[mink]}] // N
 ]
```

In[132]:=

```
pmono = (2 NN) ^ (- (4 NN u));
```

### Numerical recursions

### Figure 1B

Substitutions for sf and sm to fix the mean strength of selection across sexes ss and the ratio $a$

In[156]:=

```
Solve[{sf / sm == a, (sf + sm) / 2 == ss}, {sf, sm}]
```

Out[156]=

$$\left\{\left\{\text{sf}\to\frac{2\text{ a ss}}{1+\text{a}},\ \text{sm}\to\frac{2\text{ ss}}{1+\text{a}}\right\}\right\}$$

In[157]:=

```
ListPlot[
 {Table[{HF, cumSelSANum[Flatten[{{sf → (2 a ss)/(1 + a), sm → (2 ss)/(1 + a)} /. ss → 0.01 /. a → 2,
       hf → HF, hm → 1 - HF, ϵm → 1, ϵf → 1, rm → 0.5, rf → 0.5}],
     0.0005 * 2 * 10 000, 10 000, 50 000]}, {HF, 0.05, 0.95, 0.15}],
  Table[{HF, cumSelSANum[Flatten[{{sf → (2 a ss)/(1 + a), sm → (2 ss)/(1 + a)} /. ss → 0.01 /. a → 4,
```

```
        hf → HF, hm → 1 - HF, ϵm → 1, ϵf → 1, rm → 0.5, rf → 0.5}],
      0.0005 * 2 * 10 000, 10 000, 50 000]}, {HF, 0.05, 0.95, 0.15}],
   Table[{HF, cumSelSANum[Flatten[{{sf → (2 a ss)/(1 + a), sm → (2 ss)/(1 + a)} /. ss → 0.01 /. a → 8,
        hf → HF, hm → 1 - HF, ϵm → 1, ϵf → 1, rm → 0.5, rf → 0.5}],
      0.0005 * 2 * 10 000, 10 000, 50 000]}, {HF, 0.05, 0.95, 0.15}],
   Table[
    {H, cumSelUBNum[{s → 0.01, h → H, ϵ → 1, r → 0.5}, 0.0005 * 2 * 10 000, 50 000]},
    {H, 0.05, 0.95, 0.15}]}, PlotRange → Full, PlotStyle →
   {{PointSize[0.025], Darker[Blue]}, {PointSize[0.025], Lighter[Blue, 0.6]},
    {PointSize[0.025], Lighter[Purple]}, {PointSize[0.025], Red}}];


Plot[{cumSelSA /. hstar → h (1 - pm0 ϵ) /. ϵ → 1 /. pm0 → 0.0005 /.
     Flatten[{{sf → (2 a ss)/(1 + a), sm → (2 ss)/(1 + a)} /. ss → 0.01 /. a → 2,
       hf → HF, hm → 1 - HF, ϵm → 1, ϵf → 1, rm → 0.5, rf → 0.5}],
   cumSelSA /. hstar → h (1 - pm0 ϵ) /. ϵ → 1 /. pm0 → 0.0005 /.
     Flatten[{{sf → (2 a ss)/(1 + a), sm → (2 ss)/(1 + a)} /. ss → 0.01 /. a → 4,
       hf → HF, hm → 1 - HF, ϵm → 1, ϵf → 1, rm → 0.5, rf → 0.5}],
   cumSelSA /. hstar → h (1 - pm0 ϵ) /. ϵ → 1 /. pm0 → 0.0005 /.
     Flatten[{{sf → (2 a ss)/(1 + a), sm → (2 ss)/(1 + a)} /. ss → 0.01 /. a → 8,
       hf → HF, hm → 1 - HF, ϵm → 1, ϵf → 1, rm → 0.5, rf → 0.5}],
   (1 / 2) s h ϵ (totHetUB) /. hstar → h (1 - pm0 ϵ) /. ϵ → 1 /. pm0 → 0.0005 /. h → HF},
  {HF, 0.01, 0.99}, PlotRange → Full,
  PlotStyle → {{Darker[Blue]}, {Lighter[Blue, 0.6]}, {Lighter[Purple]}, {Red}},
  AxesOrigin → {0, 0}];

plt1B = Legended[Labeled[Show[{%, %%}, AxesStyle → Thick,
     TicksStyle → Directive[Black, 14], AxesOrigin → {0, 0}, ImageSize → 350],
   {Text[Style[Column[{"dominance in females", "(h_f)"}, Center], 16]],
     Rotate[Text[Style[Column[{"cumulative selection on", "modifier allele\n"},
         Center], 16]], π / 2], Text[Style["B", 28]]}, {Bottom, Left, {Top, Left}}],
  SwatchLegend[{Darker[Blue], Lighter[Blue, 0.6], Lighter[Purple], Red},
   fixedLabel /@ {"s_f/s_m = 2", "s_f/s_m = 4",
     "s_f/s_m = 8", "s = 0.01 (uncond. ben.)"}]]
```

Out[159]=

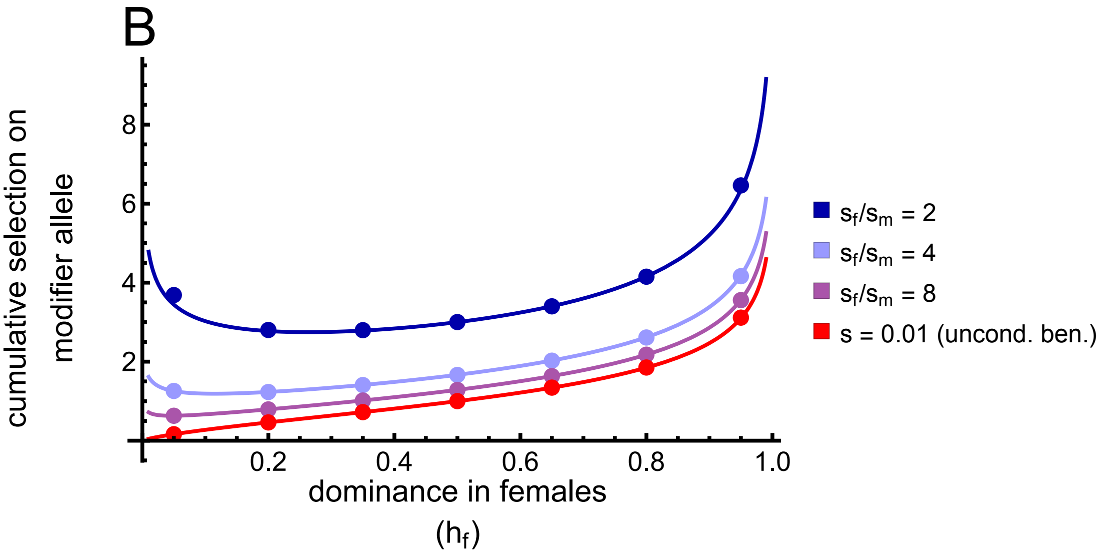

B
cumulative selection on modifier allele
8
6
4
2
0.2
0.4
0.6
0.8
1.0
dominance in females ($h_f$)
$s_f/s_m$ = 2
$s_f/s_m$ = 4
$s_f/s_m$ = 8
s = 0.01 (uncond. ben.)

### Figure 2A

In[160]:=

```
traj1 =
 Labeled[Show[{With[{data = freqListSA[{sf → 0.02, sm → 0.01, ϵf → 0.5, ϵm → 0.5,
        rf → 0.5, rm → 0.5, hf → 0.5, hm → 0.5}, 100, 1000, 8000]},

     ListPlot[{1 - data⟦All, 1⟧, 1 - data⟦All, 2⟧}, AxesStyle → Thick, PlotMarkers →
       {Automatic, Scaled[0.015]}, TicksStyle → Directive["Label", 12],
      ImageSize → 350, PlotStyle → {Blue, Orange}, PlotRange → {0, 1.01}]

    ],
    Graphics[{Darker[Green], Dashed, Thick,
      Line[{{15 000, (sf - sm)/(sf ϵf + sm ϵm) /. {sm → 0.01, sf → 0.02, hf → 0.5, hm → 0.5,
            ϵm → 0.5, ϵf → 0.5, rf → 0.5, rm → 0.5}}, {0, (sf - sm)/(sf ϵf + sm ϵm) /. {sm → 0.01,
            sf → 0.02, hf → 0.5, hm → 0.5, ϵm → 0.5, ϵf → 0.5, rf → 0.5, rm → 0.5}}}]}]
   }],
  {Style[Text["generation"], 16], Rotate[Style[Text["frequency"], 16], Pi / 2],
   Style[Text["A"], 28]}, {Bottom, Left, {Top, Left}}]
```

Out[160]=

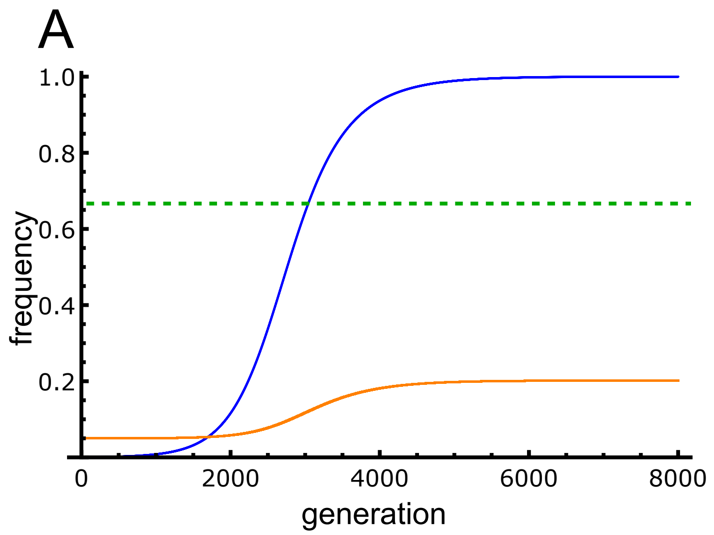

### Figure 2B

In[161]:=

```
traj2 = Labeled[
  Show[{With[{data = freqListSA[{sf → 0.02, sm → 0.01, ϵf → 1, ϵm → 1, rf → 0.5,
        rm → 0.5, hf → 0.5, hm → 0.5}, 100, 1000, 8000]},

     ListPlot[{1 - data[[All, 1]], 1 - data[[All, 2]]},
      PlotMarkers → {Automatic, Scaled[0.015]}, AxesStyle → Thick,
      ImageSize → 350, TicksStyle → Directive["Label", 12],
      PlotStyle → {Blue, Orange}, PlotRange → {0, 1.01}]],
    Graphics[{Darker[Green], Dashed, Thick, Line[
       {{20000, (sf - sm)/(sf ϵf + sm ϵm) /. {sm → 0.01, sf → 0.015, hf → 0.5, hm → 0.5, ϵm → 0.5,
            ϵf → 0.5, rf → 0.5, rm → 0.5}}, {0, (sf - sm)/(sf ϵf + sm ϵm) /. {sm → 0.01, sf → 0.015,
            hf → 0.5, hm → 0.5, ϵm → 0.5, ϵf → 0.5, rf → 0.5, rm → 0.5}}}]}]}

  ],
  {Style[Text["generation"], 16], Rotate[Style[Text["frequency"], 16], Pi / 2],
   Style[Text["B"], 28]}, {Bottom, Left, {Top, Left}}]
```

Out[161]=

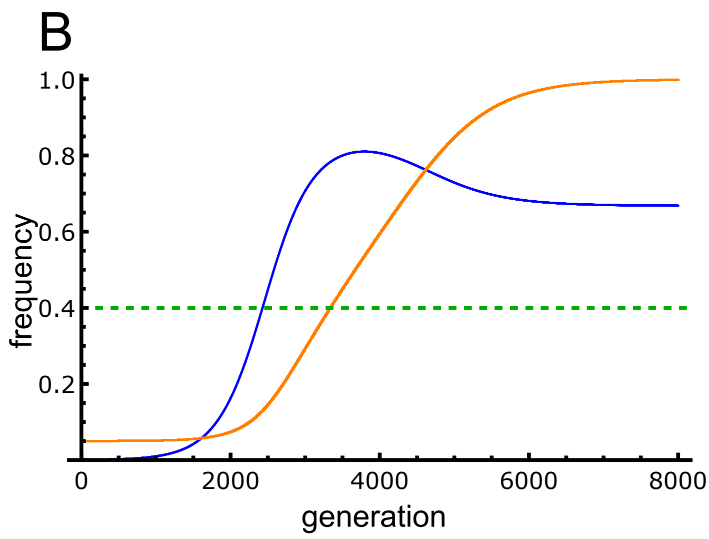


### Figure 2C

In[163]:=

```
hmapdata = Flatten[
   Table[
    {SM, SF, Last[pmListSA[{sf → SF, sm → SM, hf → 0.5, hm → 0.5, ϵm → 0.5,
         ϵf → 0.5, rf → 0.5, rm → 0.5}, 0.05 * 2 * 10 000, 10 000, 50 000]]},
    {SF, 0, 0.1, 0.0025}, {SM, 0, 0.1, 0.0025}], 1];
```

In[164]:=

```
hmap = ListContourPlot[hmapdata(*ColorFunction→Function[{x},
      If[0<x≤0.1, Gray, If[0.1<x<0.99, Lighter[Blue,0.4], Lighter[Red,0.6]]]]*),
   AxesLabel → Automatic, InterpolationOrder → 0, FrameTicksStyle → Directive[12],
   FrameStyle → Thick, PlotLegends → BarLegend["LakeColors"],
   ColorFunction → "LakeColors", ColorFunctionScaling → False, ImageSize → 380];

hmapBoundaries = Show[{hmap,
    RegionPlot[(polyConditionsMod /. sm → SM /. sf → SF /. hm → 1 / 2 /. hf → 1 / 2 /.
         ϵf → 1 / 2 /. ϵm → 1 / 2 /. pm → 0), {SM, 0, 0.1},
     {SF, 0, 0.1}, PlotStyle → None, BoundaryStyle → {Gray, Dashed}],
    RegionPlot[
     (polyConditionsMod /. sm → SM /. sf → SF /. hm → 1 / 2 /. hf → 1 / 2 /. ϵf → 1 / 2 /.
         ϵm → 1 / 2 /. pm → 0.05) || ((SF > SM) &&
        ((Chop[finalModFreqSA /. sm → SM /. sf → SF /. hm → 0.5 /. hf → 0.5 /. ϵf →
                0.5 /. ϵm → 0.5 /. pm0 → 0.05]) > (SF - SM) / (SF 0.5 + SM 0.5))),
     {SM, 0, 0.1}, {SF, 0, 0.1}, PlotStyle → None, BoundaryStyle → {Orange}],
    RegionPlot[(polyConditionsMod /. sm → SM /. sf → SF /. hm → 1 / 2 /. hf → 1 / 2 /.
         ϵf → 1 / 2 /. ϵm → 1 / 2 /. pm → 0.05), {SM, 0, 0.1}, {SF, 0, 0.1},
     PlotStyle → None, BoundaryStyle → {Thick, Black, Dashed}]}];

hmapLabeled = Labeled[hmapBoundaries, {Style[Text["sm"], 18],
    Style[Text["sf"], 18], Style[Text["C"], 28]}, { Bottom, Left, {Top, Left}}];


plt2C = Legended[hmapLabeled /.
   {_EdgeForm, c_?ColorQ, rest__} :→ {EdgeForm[Directive[Thin, c]], c, rest},
  BarLegend["LakeColors", LabelStyle → Directive["Label", 12],
   LegendLabel → Style[Text["p∞m"], 18]]]
```

Out[167]=

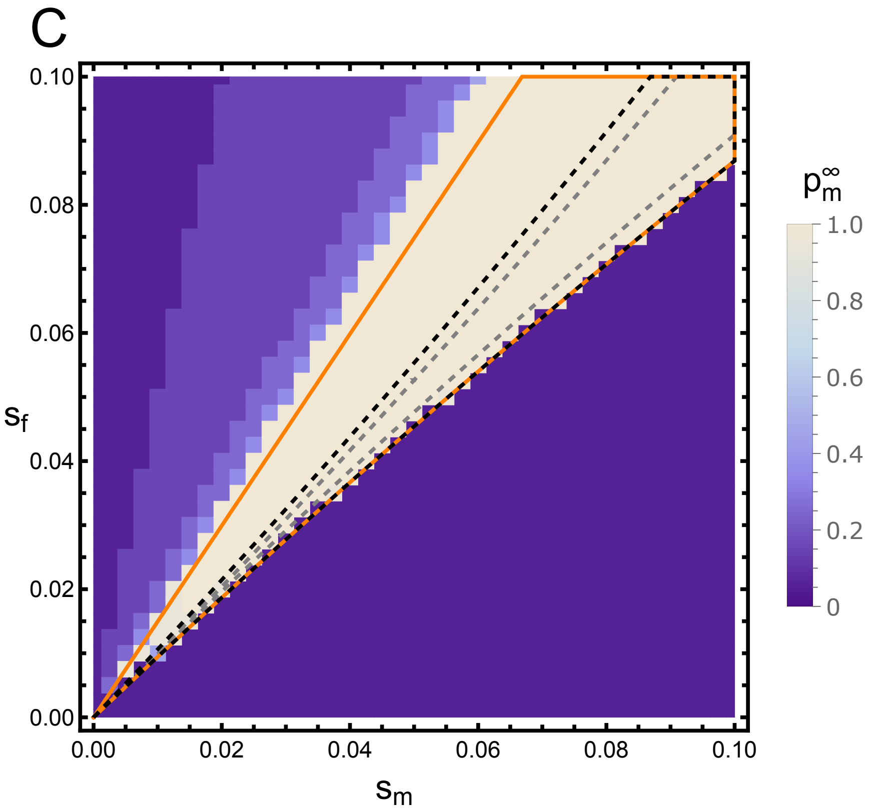
C
0.10
0.08
0.06
0.04
0.02
0.00
$s_f$
0.00
0.02
0.04
0.06
0.08
0.10
$s_m$
$p_m^\infty$
1.0
0.8
0.6
0.4
0.2
0

### Figure 1

In[168]:=

```
Style[Grid[{{plt1A}, {plt1B}}], ImageSizeMultipliers → {1, 1}]
```

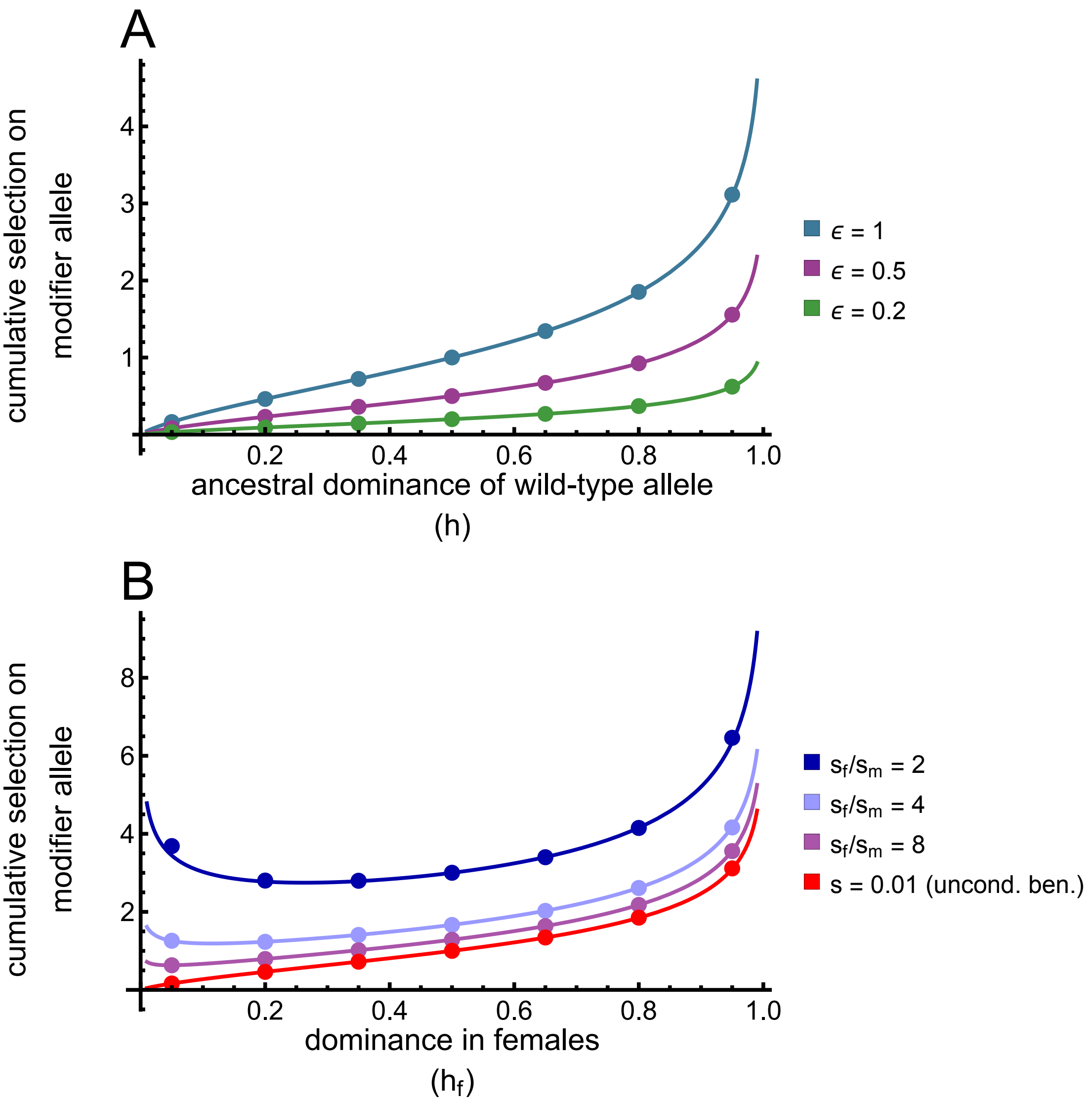

### Figure 2

In[169]:=

```
Style[Grid[{{Grid[{{traj1}, {traj2}}], plt2C}}], ImageSizeMultipliers → {1, 1}]
```

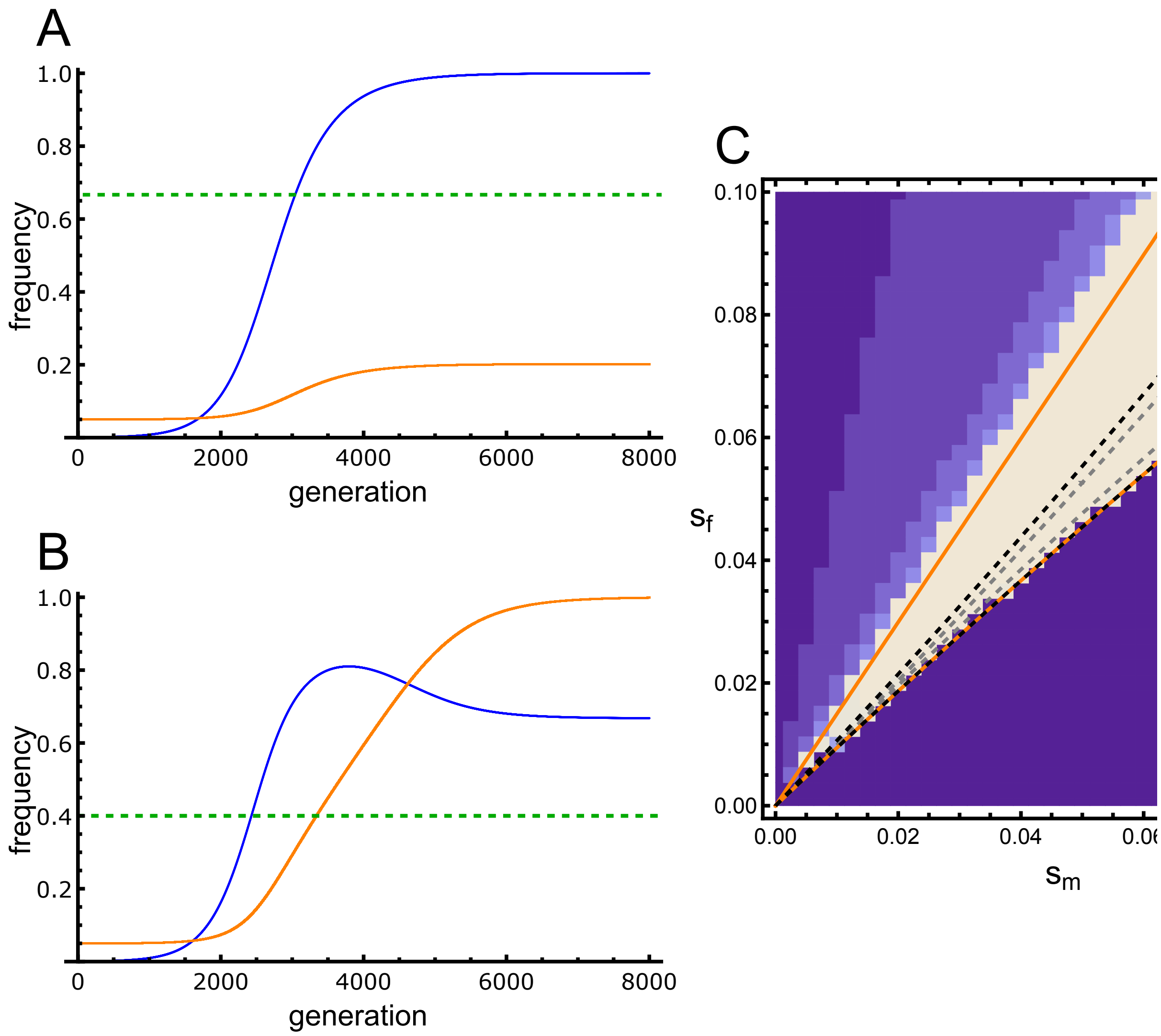

### Figure 3

In[170]:=

```
nsweepsplt =
  Quiet[ListPlot[{Table[{HF, 1/ (((1 - pmono) /. NN → 10 000 /. u → 10^-8) *
        probConversion[Flatten[{{sf → 2 a ss/(1 + a), sm → 2 ss/(1 + a)} /. ss → 0.01 /. a → 2,
           hf → HF, hm → 1 - HF, ϵm → 1, ϵf → 1, rm → 0.5, rf → 0.5}],
          10 000])}, {HF, 0.01, 0.99, 0.01}],
     Table[
      {HF, 1/ (((1 - pmono) /. NN → 10 000 /. u → 10^-8) * probConversion[Flatten[
             {{sf → 2 a ss/(1 + a), sm → 2 ss/(1 + a)} /. ss → 0.01 /. a → 4, hf → HF, hm → 1 - HF, ϵm →
               1, ϵf → 1, rm → 0.5, rf → 0.5}], 10 000])}, {HF, 0.01, 0.99, 0.01}],
     Table[{HF, 1/ (((1 - pmono) /. NN → 10 000 /. u → 10^-8) *
          probConversion[Flatten[{{sf → 2 a ss/(1 + a), sm → 2 ss/(1 + a)} /. ss → 0.01 /. a → 8,
             hf → HF, hm → 1 - HF, ϵm → 1, ϵf → 1, rm → 0.5, rf → 0.5}], 10 000])},
      {HF, 0.01, 0.99, 0.01}]}, Joined → True, PlotRange → {0, 2500},
    AxesStyle → Thick, TicksStyle → Directive[Black, 12],
    PlotStyle → {{Darker[Blue]},
      {Lighter[Blue, 0.6]}, {Lighter[Purple]}, {Red}}]];
```

In[175]:=

```
nsweepsLabeled = hmapLabeled =
   Labeled[nsweepsplt, {Rotate[Style[Text["# sweeps"], 18], π / 2], Style[Text[
       "dominance in females \n                    (hf)"], 18]}, { Left, Bottom}];
```

In[176]:=

```
fig3 = Legended[nsweepsLabeled,
  SwatchLegend[{Darker[Blue], Lighter[Blue, 0.6], Lighter[Purple]},
   fixedLabel /@ {"sf/sm = 2", "sf/sm = 4", "sf/sm = 8"}]]
```

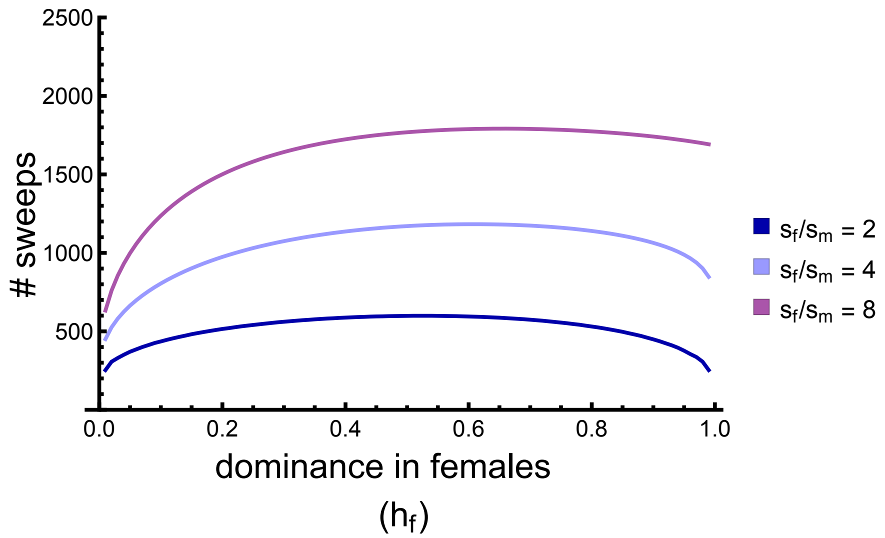
